# Patterns of numtogenesis in sixteen different mice strains

**DOI:** 10.1101/2022.08.05.502930

**Authors:** Bálint Biró, Zoltán Gál, Michael Brookman, Orsolya Ivett Hoffmann

## Abstract

Numtogenesis is the phenomenon of mitochondrial sequence localisation and integration into the nuclear genome. This is an ongoing process which contributed to the complexity of eukaryotic genomes. The sequences that are integrated into the nuclear genome are called nuclear mitochondrial sequences (numt). numts have a wide variety of applications in tumor biology, phylogenetic studies, forensic research and so on. *Mus musculus musculus* is the most popular model organism. Numerous mouse strains are used in medical research to model human diseases. Numts were described in the genome of *Mus musculus musculus* just like in many other species however the characterisation of numts in different mouse strains is missing. In this study we explored the patterns of numtogenesis in 16 mouse strains by aligning the nuclear genomes with the corresponding mitochondria. Investigation of numts shed light on strain specific differences and resembles the phylogenetic relationships as to our current knowledge in most of the cases.

## 2 Introduction

### 2.1 Definition and importance of numtogenesis

The sequences in the nuclear genome with mitochondrial origins are called numts (Lopez et al., 1994) and their integration process itself is called numtogenesis (Singh et al., 2017).

The importance of numts were proved in several pathophysiological conditions where oncogene activation, tumor supressor deactivation were the consequences of numt insertion (Singh et al., 2017; Srinivasainagendra et al., 2017; Palodhi et al., 2020). Other than cancer biology, numts have important applications in the fields of phylogenetics (Ko et al., 2015; Nacer & do Amaral, 2017) and forensic research (Marshall & Parson, 2021; Cortes-Figueiredo et al., 2021). Based on recent studies, numts have an influence on mitochondrial genome editing systems by providing OFF-target cleavage sites in the nuclear genome (Lei et al., 2022).

### 2.2 Molecular background of numtogenesis

The precondition of numtogenesis is the degradation of the mitochondrial membrane which is due to mitostress inducing factors (Gaziev & Shaikhaev, 2007). These factors could be ionizing radiation, endotoxins, ROS, aging, heat shock etc.. After the membrane damage has happened, the organelle is going to be tagged with a degradation signal which will induce mitophagy. Mitophagy is a special case of autophagy which takes place in the mitophagosome (Goldman et al., 2010). Mitophagosome is responsible for eliminating and recycling damaged mitochondria (Hazkani-Covo et al., 2010). During imperfect mitophagy, mitochondrial components (i.e. numts) could escape into the cytoplasm. Another explanation is that numts could enter into the cytosol during fusion and fission events (Hazkani-Covo et al., 2010; Puertas & González-Sánchez, 2020). Numts are protected in the cytoplasm against nucleases due to a vacuole mediated process or due to complex formation with histone like proteins. These cytosol located numts can enter the nucleus by membrane fusion (Gaziev & Shaikhaev, 2007). Once the numts are in the nucleus, the Non-Homologous-End-Joining repair mechanism (NHEJ) will insert them into the nuclear genome at double stranded brakes. In the case of NHEJ when no template DNA is available, a nuclease mediated deletion is likely to happen. This will result in a relatively long, single stranded DNA which risks the occurrences of long deletions and translocations. According to alternative explanations, numts serve as template DNAs to prevent bigger mutations (Hazkani-Covo et al., 2010).

### 2.3 Evolutionary background of numtogenesis

At the very beginning of the eukaryotic evolution, an intracellular symbiotic partnership happened between proteobacteria and Archea (W. F. Martin et al., 2015; Roger et al., 2017). Throughout this long specialisation process, different intracellular organelles were formed to execute different functions. One of these cellular compartments is mitochondrion which serves as the main machinery for aerobic respiration and participates in many other important cellular processes (Roger et al., 2017). Mitochondrion also has its own genome which was seriously reduced in size through the coevolution of symbiont and host. This reduction process is called endosymbiotic gene transfer (EGT) and basically it is a genome plasticity phenomenon since parts of mitochondrion’s genetic material are integrated into the nuclear genome (Kelly, 2020). Muller’s theory (aka Muller’s ratchet) serves as an explanation to EGT which declares that deleterious mutations are going to reduce the size of an asexually isolated genome. Hence in long term the genome will go extinct (Muller, 1964; Metzger & Eule, 2013; Naito & Pawlowska, 2016). The uniparental inheritance of the mitochondrial genome asexually isolates it since it cannot be recombined (Breton & Stewart, 2015). This means that Muller’s ratchet is applicable to the mitochondrial genome (Howe & Denver, 2008). Therefore EGT prevents mtDNA to be totally degraded as it could happen as proposed by Muller by inserting numts into the nuclear genome (W. Martin & Herrmann, 1998; Kelly, 2020). Another explanation to EGT is that it is too costly for a cell to maintain a highly complex interactome network of multiple organelles and multiple genomes per each organelle. That is why EGT is functioning (Kelly, 2021).

### 2.4 Objective

Mouse is the most commonly used model organism and numerous mice strains serve as model animals for important human diseases (Perlman, 2016; Hickman et al., 2017; Tibbetts, 2018; Murillo-Cuesta et al., 2020). Numts have been characterized in several species like honeybee (Behura, 2007), human (Simone et al., 2011), wasps (Wang et al., 2020), cat (Lopez et al., 1994) and so on. Numts were also described even in mouse genome (Calabrese et al., 2012) but no detailed characterisation were published in terms of different mice strains. Therefore our primary goal was to explore the patterns of numts in different mice strains and to see whether they are different from the numts in *Mus musculus musculus*.

## 3 Materials and Methods

Nuclear and mitochondrial DNAs were acquired from Ensembl and NCBI databases (Wheeler et al., 2007; Cunningham et al., 2022). The mitochondrion of *Mus musculus musculus* was annotated using MITOS server hosted by the University of Leipzig (Bernt et al., 2013). For the numt comparisons 12 inbred strains (*129s1svimj, aj, akrj, balbcj, c3hhej, c57bl6nj, cbaj, dba2j, fvbnj, lpj, nzohiltj, and nodshiltj*) and four ”wild derived” strains (*casteij, pwkphj, wsbeij, spreteij*) were selected based on a previously published genome comparison analysis (Lilue et al., 2018). Double mtDNA and gDNA were aligned using LASTAL (v1219) (Kielbasa et al., 2011) with the scoring scheme of + 1 for matches, −1 for mismatches, 7 for gap-open penalty and 1 for gap-extension penalty. Using doubled mtDNA, we were able to identify numts that are located on the linearization point of the mtDNA. After that, alignments were filtered based on their e-values as proposed by Tsuji (Tsuji et al., 2012). False positive alignments that were the results of using double mtDNA were also discarded.

To examine normality, Anderson-Darling test was performed at 0.05 p value. t-test or Wilcoxon signed rank test was performed based on the result of normality testing. All the statistical calculations were conducted in Scipy (v1.6.2) (Virtanen et al., 2020). Whether tRNA genes are expressed were derived from their structures using free energy values which were calculated with seqfold (Ouyang et al., 2013) based on a previous research paper (Liu & Zhao, 2007).

Pairwise divergences were calculated using the modified Kimura 2 parameter (Nishimaki & Sato, 2019) which tolerates gaps in the alignments. The modified Kimura 2 formula is described by Equation 3.1.

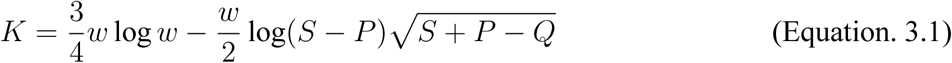

where *w*=the probability that the given position contains a nucleotide,

*S*=*n*_1_/*n*,

*n*_1_=the number of positions where the two aligned sequences contain the same nucleotide,

*n*=total number of nucleotides,

*P* =*n*_2_/*n*,

*n*_2_=number of transition type mutations,

*n*_3_=number of transversion type mutations.

Pairwise divergence values were calculated using a sliding window approach with 1kb window size and 10bp step size.

For the free energy calculations, the corresponding tRNA sequences for each strains were acquired with SAMTOOLS’s (v1.6) faidx function (Li et al., 2009).

The phylogenetic analysis was conducted in R with phangorn (Schliep, 2011). The Maximum-Parsimony tree was constructed using Jukes-Cantor distance with 100 bootstrap cycles. For tree construction cytochrome b (*CYTB*) and corresponding numt sequences were used as described by a previous study (Rodríguez et al., 2007) with rat as outgroup.

All figures were created in matplotlib (v3.4.3) and seaborn (v0.11.2) (Hunter, 2007; Waskom, 2021).

Plain LaTex file and analysis codes are available at https://github.com/balintbiro/Patterns-of-numtogenesis-in-sixteen-different-mice-strains GitHub repository.

## 4 Results

152 numts can be identified in the *Mus musculus musculus* genome. There is a huge variability in terms of numt numbers on the chromosomes. For example, chr1 contains 14 numts, while no numt can be found on chr19. The numts cover the total mitochondrion. The longest numt (4654 bp) covers 6 protein coding genes (Fig.1/a). There is a strong correlation (Pearson correlation coefficients: 0.77) between the chromosome size and the number of numts on a given chromosome (Fig.1/b). The shortest chromosome contains the smallest number of numts while the longest chromosome contains the highest number of numts. However this relationship is not true for any of the strains investigated.

**Fig. 1.**
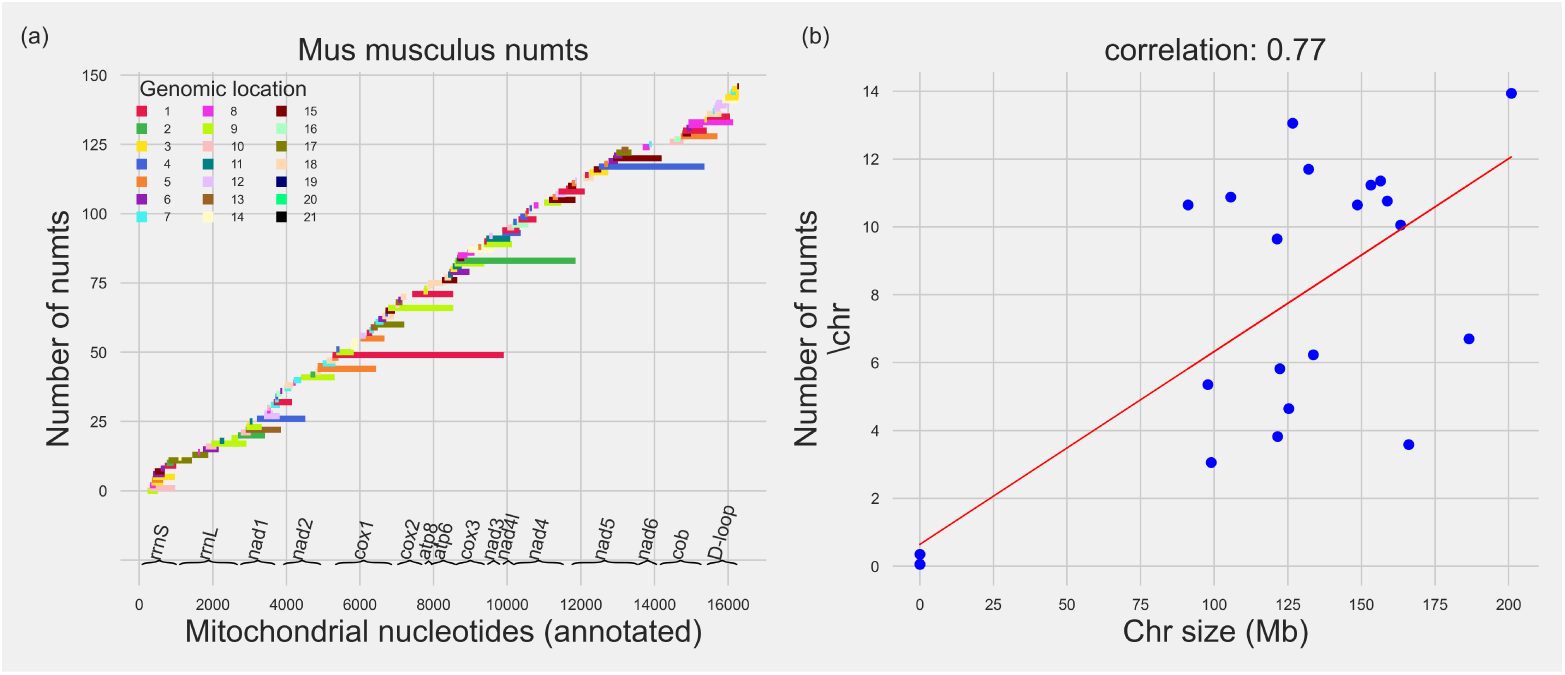
Patterns of *Mus musculus musculus* numts. Distribution of *Mus musculus musculus* with the genomic locations (a) and the correlation between the number of numts and the size of the corresponding chromosome (b). Small tRNA genes are not part of the annotation.

When investigating the distribution of numts along the mitochondria in different strains, it turns out that the majority of the numts are the same as in the case of *Mus musculus musculus*. However exceptions do exist. For example as we have already described above, numts originate from the whole mitochondrion in case of *Mus musculus musculus* but not in the case of several inbred and wild derived strains. Differences are also present in the lengths of the numts. For instance, the longest numt in the *Mus musculus musculus* genome which locates on chr1 is missing from all the strains investigated (Fig.2.).

**Fig. 2.**
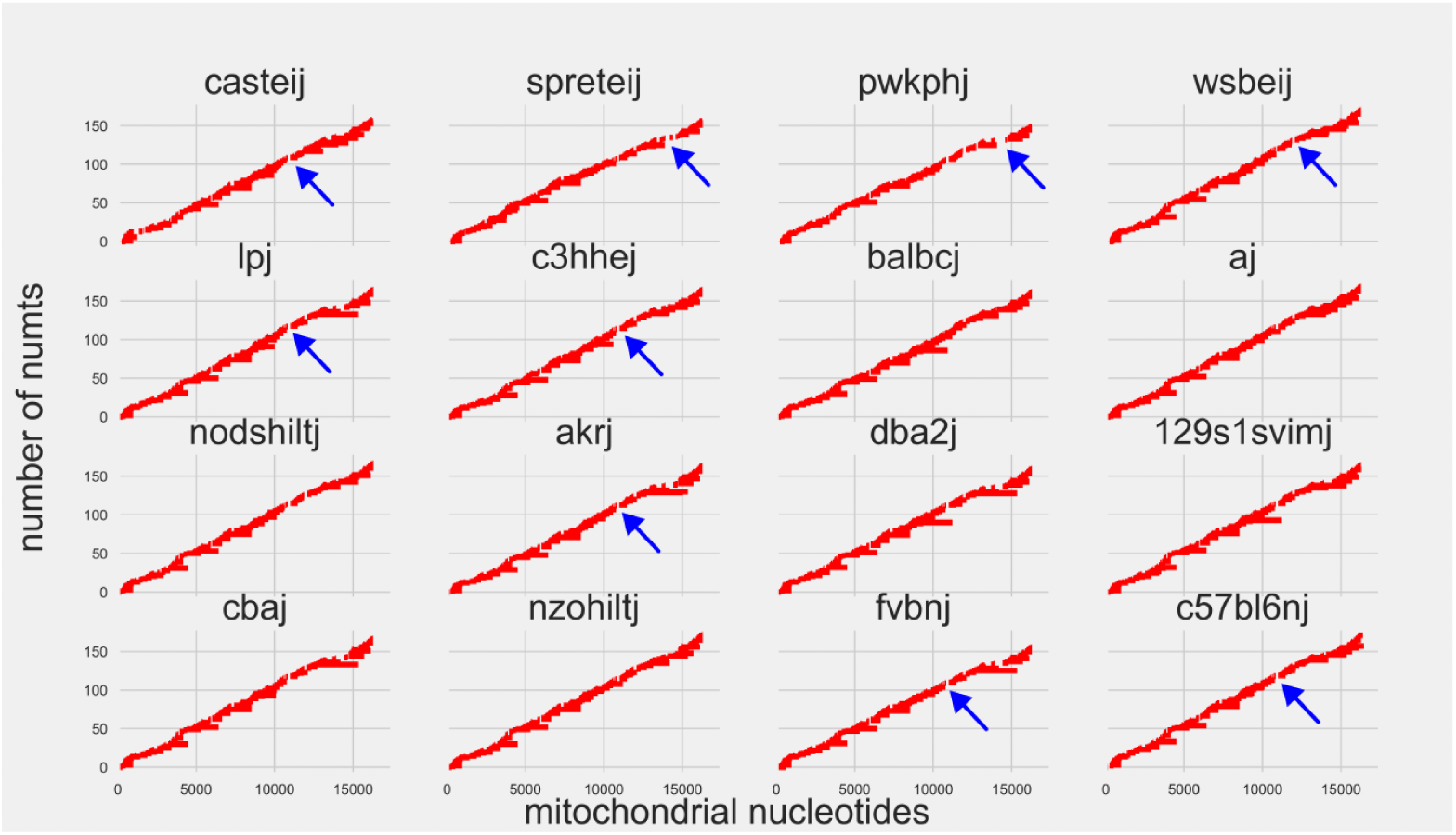
Mitochondrial origin of numts in different mice strains. The upper row contains the wild derived strains. Blue arrows indicate mitochondrial regions where numts do not originate from.

There are common patterns in terms of numt coverage. Interestingly the regions of mitochondria that contain the linearization point (the start and the end of the linearized mitochondrion) are highly covered by numts. This region contains the *tRNA-S* coding gene and the D-loop. Another numt dense region which is present in every strains, covers *atp6, cox3, nad3* and *nad4l* genes. There are two general numt sparse regions. The first one partially covers *tRNA-S* and *tRNA-L* genes while the second one covers *nad4* (Fig.3.).

**Fig. 3.**
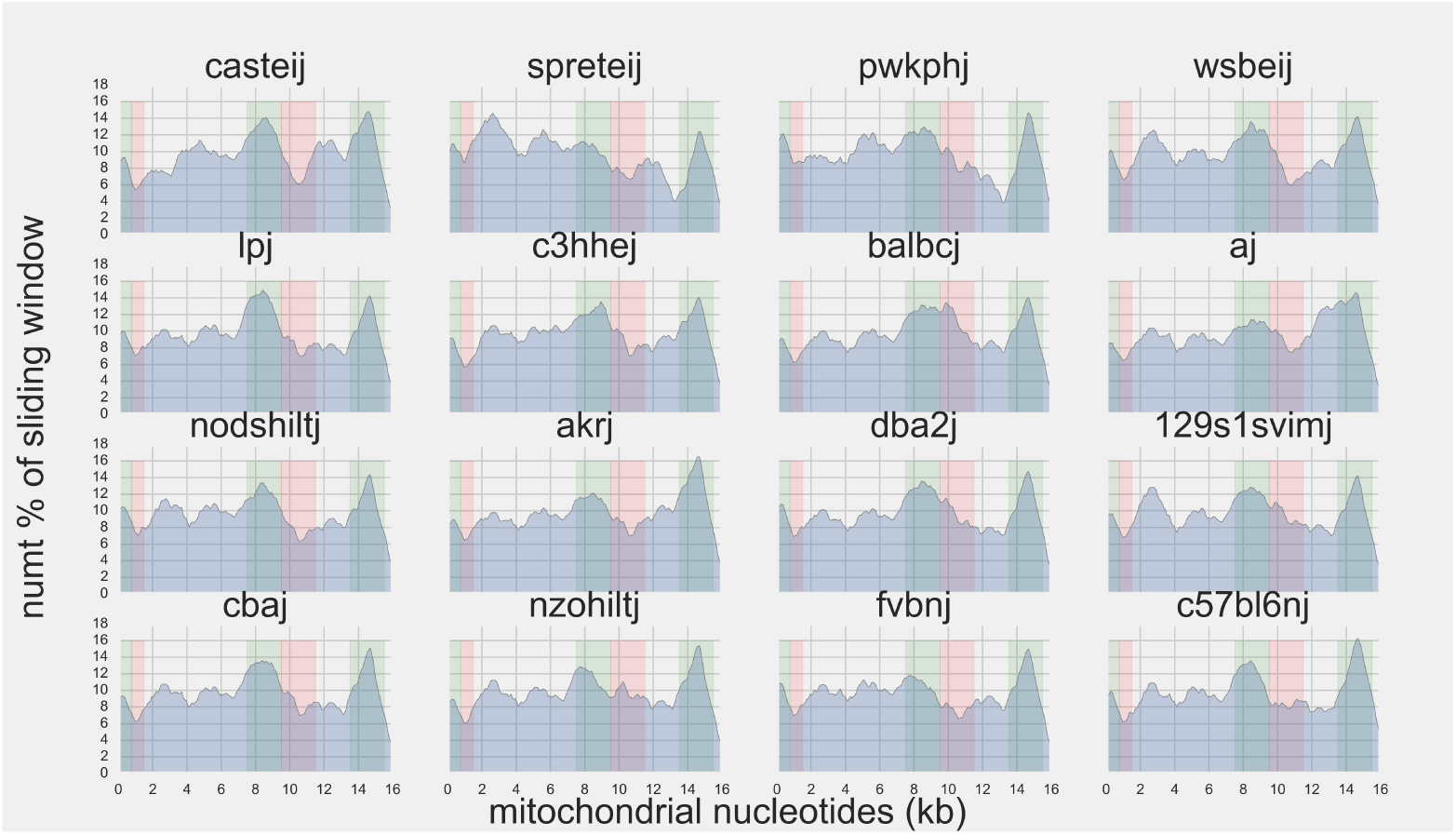
Numt contents along mitochondria. Colored areas show common patterns. Green coloring is corresponding for dense numt region while red coloring is corresponding for numt sparse region.

From the wild derived strains *casteij, spreteij* and *pwkphj* while from inbred strains *aj* and *akrj* are clustered together when it comes to nuclotides along mitochondria involved in numtogenesis. At the same time *wsbeij* and the rest of the inbred strains are also clustered together (Fig.4.).

**Fig. 4.**
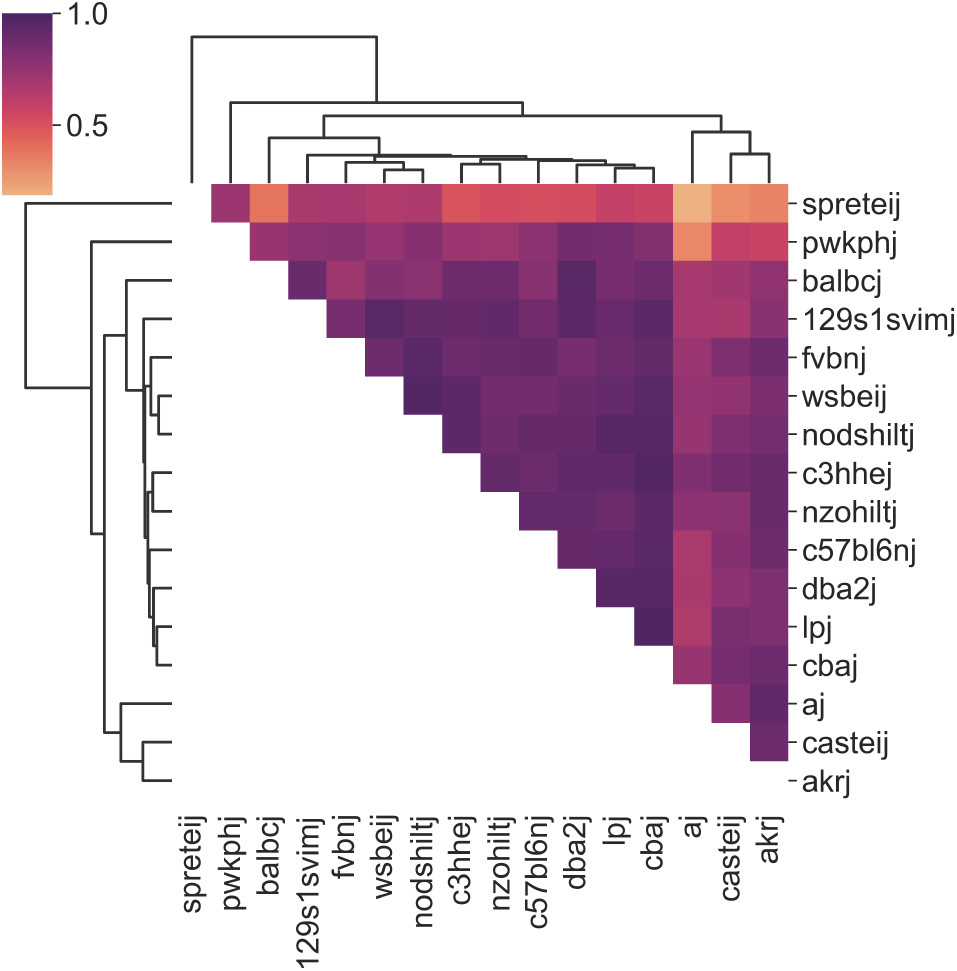
Clustered heatmap representation of the Pearson correlation matrix of nuclotides involved in numtogenesis along mitochondria.

The wild derived strains *casteij, spreteij* and *pwkphj* show higher pairwise divergence values when total mitochondria are compared with the mitochondrion of *Mus musculus musculus*. Surprisingly, the fourth wild derived strain *wsbeij* does not differ significantly from *Mus musculus musculus* while the inbred *nzohiltj* strain shows elevated pairvise distance values to some extent (Fig.5.).

**Fig. 5.**
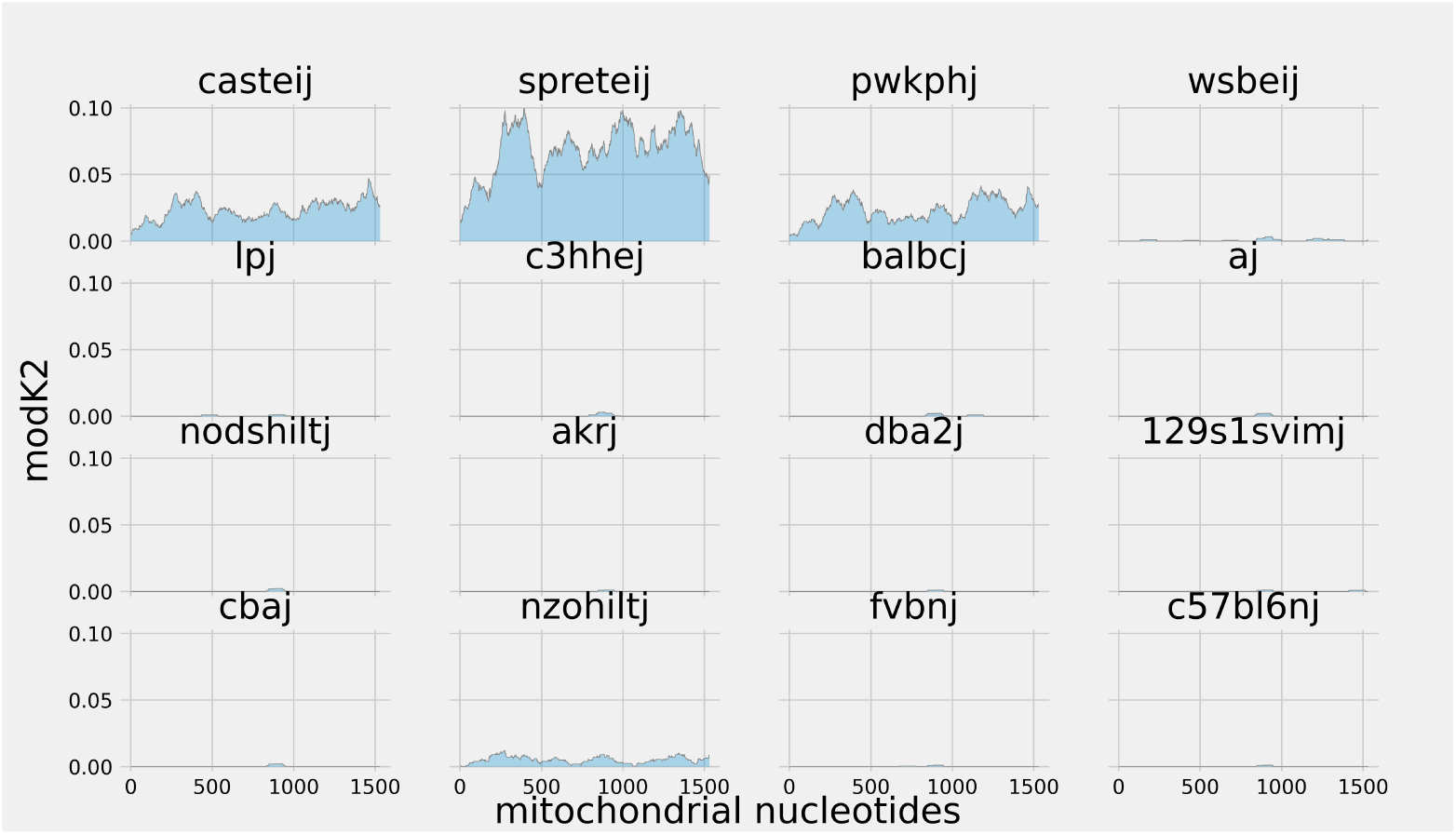
Pairwise divergence values. The upper row contains the wild derived strains.

Maxmimum-Parsimony phylogenetic tree supports two monophyletic clades, namely the mitochondrial CYTB and the corresponding numt sequences. The result of the phylogenetic analysis resembles the whole mitochondria divergence values even though the sequences of only one gene were used. In the CYTB clade the wild derived strains *casteij, spreteij* and *pwkphj* plus the inbred *nzohiltj* strain differ from the other strains. However *nzohiltj* is very close to the other inbred strains. In the numt clade the wild derived strains *casteij, spreteij* and *pwkphj* are different than the rest of the strains. In this clade the inbred strain *nzohiltj* is clustered together with the inbred strains unlike in the case of CYTB clade, pairwise divergence and numt coverage analysis. Intriguingly *wsbeij* is clustered together with the inbred strains in both clades even though it is a wild dervied strain (Fig.6.).

**Fig. 6.**
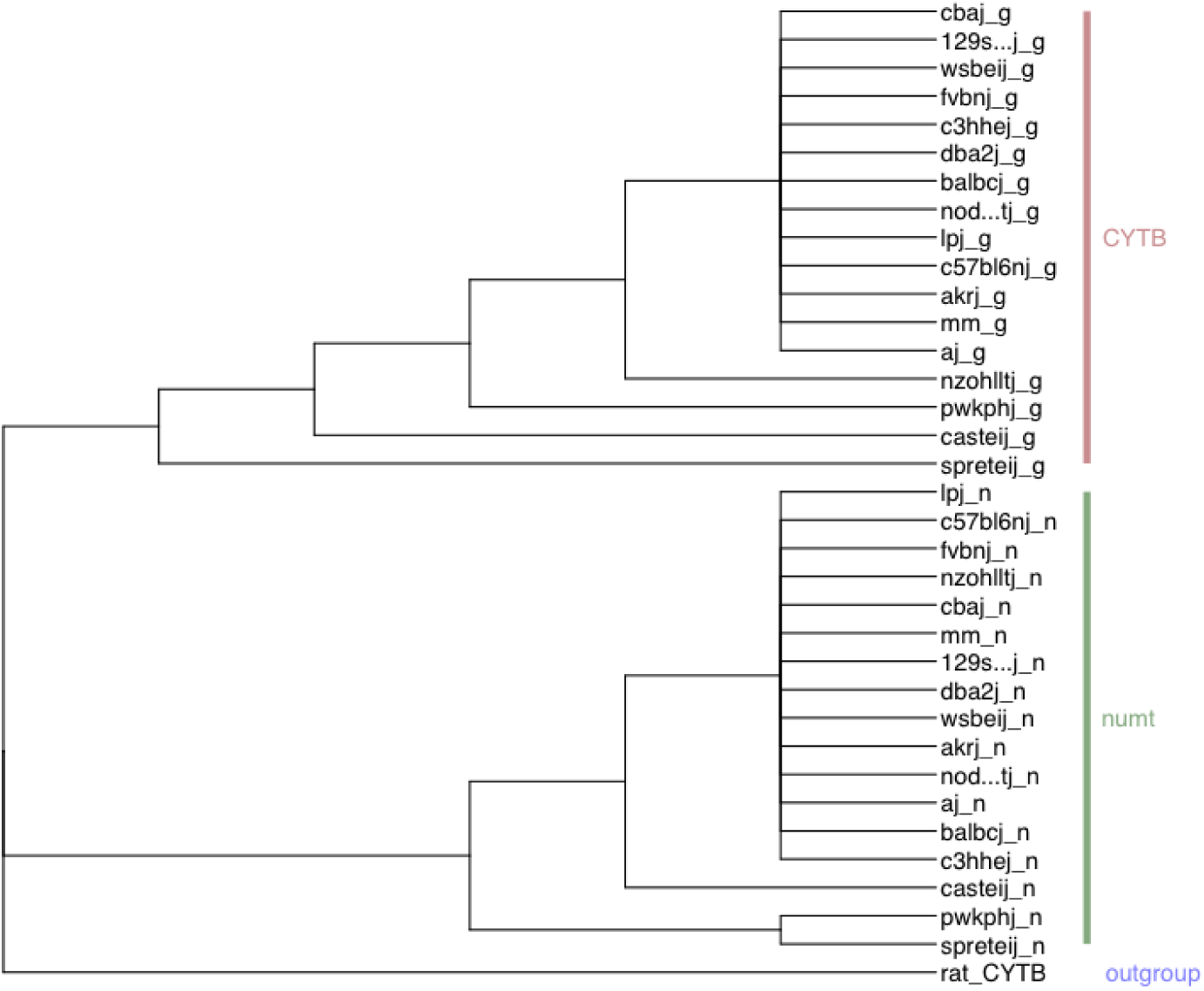
Maximum-Parsimony tree based on Jukes-Cantor distance with 100 bootstrap and rat CYTB as outgroup.

During the analysis of free energy values of the folding of tRNA sequences and their corresponding numts no case was shown where numt folding’s ΔG value was the same as tRNA folding’s. In most of the cases tRNA folding’s ΔG was smaller than numt folding’s ΔG (Fig.7.).

**Fig. 7.**
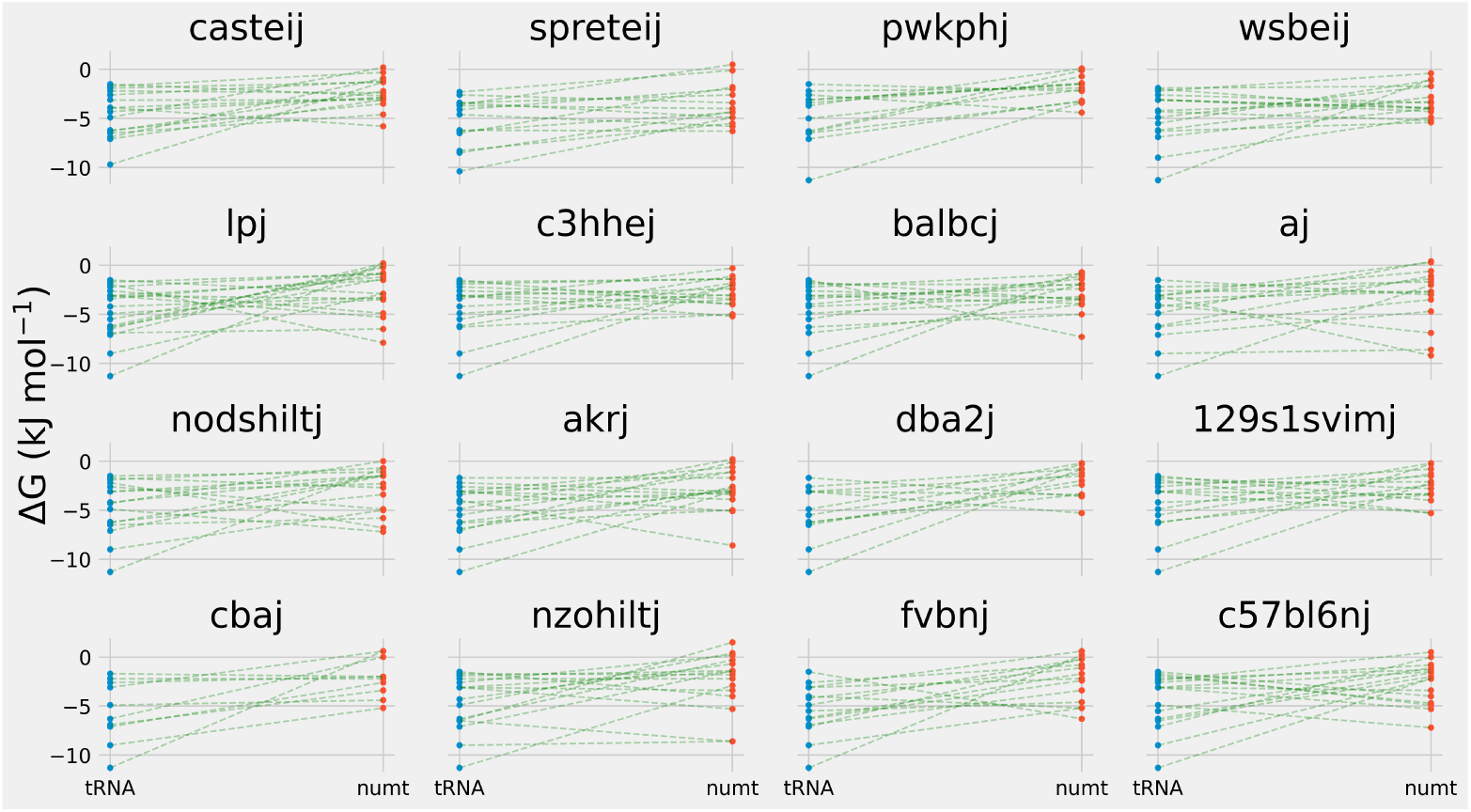
The free energy values of the predicted structures of tRNAs and their corresponding numts. The upper row contains the wild derived strains.

## 5 Conclusions

Like in many other eukaryotic organisms (Hazkani-Covo et al., 2010; Calabrese et al., 2012; Tsuji et al., 2012; Wang et al., 2020), numts are also present in the genome of *Mus musculus musculus*. However the different patterns of numts in mice strains have not been described previously. Hence in this study numts of divergent mice strains were investigated.

We described 152 numts in the *Mus musculus musculus* genome which is comparable with previous studies (Calabrese et al., 2012; Tsuji et al., 2012).

There is a strong correlation in the *Mus musculus musculus* genome between chromosomal length and the number of the numts on the given chromosome. This correlation also exists in the human genome (Lascaro et al., 2008; Simone et al., 2011). In general, there is a higher gene density in case of shorter chromosomes. Since natural selection tries to avoid insertional mutagenesis (eg. intragenic numts), there is a higher possibility for less gene dense, longer chromosomes to tolerate a numt integration without serious consequences (Lascaro et al., 2008). However this is not the case in any of the strains investigated. No correlation was found between the above mentioned attributes also in the case of honey bee (Behura, 2007).

The majority of *Mus musculus musculus* numts were the same in the strains investigated. However the longest numt (4654 bp on chr1) could not be detected in any of the strains. This situation was found in a couple of other organisms and it is originated from a fragmentation event after the given numt has been integrated into the nuclear genome (Behura, 2007; Rodríguez-Salinas et al., 2012).

The investigation of numt coverage along the mitochondria reveals that mitochondrial nucleotides are present in several copies. Numt coverage results also proved that the representation of the nucleotides along the mitochondria differ. Hence that over-as well as under-represented regions do exist. This kind of imbalanced representation of mitochondrial nucleotides were reported in several mammalian species (Simone et al., 2011; Tsuji et al., 2012).

All of our experiments that somehow integrate data about the similarity of numts or mitochondria (numt content along mitochondria, pairwise divergence, phylogeny) resemble the relationship between *Mus musculus musculus* and *spreteij* as to our current knowledge. Namely that from the strains investigated, *spreteij* seems to be the most distantly related compared to *Mus musculus musculus*. The phenomenon that *spreteij* is the most distant strain from the strains examined also applies to when total nuclear genomes are being compared (Lilue et al., 2018).

Since the mitochondrial genetic code and the nuclear genetic code are different (Gonzalez et al., 2012), the same DNA sequence would be resulted in different RNA molecules depending on the used genetic code. In addition, the nucleotide composition of numts are adjusted to the mitochondrial machinery. Hence numts are considered to be pseudogenes that are unable to code the genetic information that they used to code. However there is still a chance for numts to code functional tRNAs since tRNAS coded by the nucleus and tRNAs coded by mitochondrion would be identical (Pozzi & Dowling, 2019). Although we found no evidence for expressing numts that are corresponding to tRNA genes. All the tRNA numts were altered in sequence and the free energy of the folding of these numts were always higher than the free energy of the folding of corresponding tRNA genes. However the free energy values of the folding of numts were still negative, which means that the folding itself is a spontaneous process. But tRNA genes had the minimum free energy which means the most stable structure from a thermodynamic point of view (Su et al., 2019). Our results regarding tRNA expression are in agreement with the results in the case of cattle (Liu & Zhao, 2007).

In this paper we described the patterns of numts in different mice strains and characterised the differences between each strains. Our results can contribute to distinct phylogenetic and disease model studies.

## 6 Acknowledgements

B.B. received NTP-NFTÖ-21-B-0127 scholarship. O.I.H. was funded by NKFIH OTKA FK 124708, János Bolyai Research Scholarship of the Hungarian Academy of Sciences and New National Excellence Program of the Ministry for Innovation and Technology from the source of the National Research, Development and Innovation Fund (ÚNKP-21-5).

## Notes

https://github.com/balintbiro/Patterns-of-numtogenesis-in-sixteen-different-mice-strains

### Competing Interest Statement

The authors have declared no competing interest.

### Summary of Updates

References have been added to the original definitions of numt and numtogenesis. Furthermore some typos have been corrected.

